# A United Statistical Framework for Single Cell and Bulk Sequencing Data

**DOI:** 10.1101/206532

**Authors:** Lingxue Zhu, Jing Lei, Bernie Devlin, Kathryn Roeder

## Abstract

Recent advances in technology have enabled the measurement of RNA levels for individual cells. Compared to traditional tissue-level bulk RNA-seq data, single cell sequencing yields valuable insights about gene expression profiles for different cell types, which is potentially critical for understanding many complex human diseases. However, developing quantitative tools for such data remains challenging because of high levels of technical noise, especially the “dropout” events. A “dropout” happens when the RNA for a gene fails to be amplified prior to sequencing, producing a “false” zero in the observed data. In this paper, we propose a Unified RNA-Sequencing Model (URSM) for both single cell and bulk RNA-seq data, formulated as a hierarchical model. URSM borrows the strength from both data sources and carefully models the dropouts in single cell data, leading to a more accurate estimation of cell type specific gene expression profile. In addition, URSM naturally provides inference on the dropout entries in single cell data that need to be imputed for downstream analyses, as well as the mixing proportions of different cell types in bulk samples. We adopt an empirical Bayes approach, where parameters are estimated using the EM algorithm and approximate inference is obtained by Gibbs sampling. Simulation results illustrate that URSM outperforms existing approaches both in correcting for dropouts in single cell data, as well as in deconvolving bulk samples. We also demonstrate an application to gene expression data on fetal brains, where our model successfully imputes the dropout genes and reveals cell type specific expression patterns.

## 1. Introduction

A biological organism is made up of individual cells, which work in concert in tissues to constitute functioning organs. Biologists have long thought that the key to understanding most human diseases lies in understanding the normal and abnormal function of cells. Yet, until very recently, our view of what molecules are expressed and where and when was limited to the level of tissues. Indeed RNA sequencing (RNA-seq) was introduced as a critical tool to answer these questions, but the RNA itself was collected from tissues. This bulk RNA-seq data provides reliable measurements of gene expression levels throughout the genome for bulk samples. With sufficient sequencing depth, even weakly expressed transcripts can be accurately captured by RNA-seq data. This technology has led to breakthroughs in various fields. For example, Fromer et al. (2016) use bulk data, obtained from prefrontal cortex of post-mortem subjects, to gain insight into how genetic risk variation for schizophrenia affects gene expression and likely generates risk for this severe psychiatric disorder.

Still bulk RNA-seq data inevitably ignores the heterogeneity of individual cells because the measurements are summed over the population of cells in the tissue. Yet it is reasonable to predict that diseases like schizophrenia do not arise from malfunctioning brain tissue, per se, but rather certain malfunctioning cells within that tissue. A leading hypothesis is that schizophrenia arises from synaptic dysfunction, and synapses are fundamental to neurons, so should neurons alone be targeted for analyses into schizophrenia? Actually, brain tissue is composed of a remarkably heterogeneous set of cell types, which have vastly different functions and expression profiles. While many are different types of neurons, many others support and alter the function of those neurons and their synapses. Thus, the different gene expression profiles for distinct cell types can have profound functional consequences. These likely are critical for the development of tissues and human diseases, and will be especially important as we aspire to fix such complex diseases as schizophrenia.

It is also of interest to link gene expression with genetic variation, particularly damaging variants associated with risk of disease. Until recently researchers have assumed that most cells express both copies of a gene equally; however, new findings suggest an even more complex situation motivating single cell measurements. Apparently some neurons preferentially express the copy of a gene inherited from one parent over the other and this can shape how mutated genes are expressed at the cellular level (Huang et al., 2017a).

One approach to characterize cell type specific gene expression profiles is to perform deconvolution on bulk RNA-seq data. Consider an observed gene expression matrix *X* ∈ ℝ^*N*×*M*^ for *N* genes in *M* bulk samples, each containing *K* different cell types. The goal of deconvolution is to find two non-negative matrices *Ã* ∈ ℝ^*N*×*K*^ and *W* ∈ ℝ^*K*×*M*^, such that

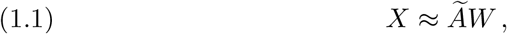

where each column of *W* represents the mixing proportion of *K* cell types in each bulk sample, and each column of *Ã* represents the average gene expression levels in each type of cells. If the “signature” matrix *Ã* is available for a set of “barcode genes” in each cell type, deconvolution reduces to a regression-type problem that aims at estimating *W*. Several algorithms have been proposed under this setting, including Cibersort (Newman et al., 2015) and csSAM (Shen-Orr et al., 2010). However, without knowing the signature matrix, deconvolution is highly nontrivial, and this line of methods includes the Deconf algorithm (Repsilber et al., 2010), semi-supervised Nonnegative Matrix Factorization algorithm (ssNMF) (Gaujoux and Seoighe, 2012), and Digital Sorting Algorithm (DSA) (Zhong et al., 2013).

A fundamental challenge of the NMF-based methods is the non-uniqueness of the factorization (Donoho and Stodden, 2003). Therefore, to obtain a biologically meaningful result, both ssNMF (Gaujoux and Seoighe, 2012) and DSA (Zhong et al., 2013) use a set of “marker genes” to guide the factorization. A marker gene is a gene that only expresses in one cell type. In other words, there are several rows of *Ã* that are priorly known to be nonzero at only one column. This is equivalent to the separability assumption introduced by Donoho and Stodden (2003) for the uniqueness of NMF. Unfortunately, marker genes are rarely known in practice. In fact, extracting high-quality marker genes is a challenging step, which is often approached by analyzing purified cells (Abbas et al., 2009).

On the other hand, single cell RNA sequencing provides gene expression measurements in individual cells, yielding a high-resolution view of cellular states that are uncharacterized in bulk data. Recent advances in high-throughput technologies have made it possible to profile hundreds and thousands of cells (Kolodziejczyk et al., 2015; Fan, Fu and Fodor, 2015). With several extra pre-processing steps including reverse transcription and amplification, the single cell mRNA library goes through similar sequencing procedures as the bulk samples, and the gene expression levels are measured by the number of mapped reads. With single cell RNA-seq data, one can investigate distinct subpopulations of cells, gain better understanding of the developmental features of different cell types (Grün et al., 2015), identify cellular differences between healthy and diseased tissues (Kharchenko, Silberstein and Scadden, 2014), and infer gene-regulatory interactions (Padovan-Merhar and Raj, 2013).

The challenges of modeling single cell RNA-seq data come from high cell-to-cell variation, as well as high levels of technical noise during sequencing due to the low amounts of starting mRNAs in individual cells. One important bias comes from the so-called “dropout” events. A dropout happens when a transcript is not detected due to failure of amplification prior to sequencing, leading to a “false” zero in the observed data (Kolodziejczyk et al., 2015). Given the excessive amount of zero observations in single cell RNA-seq data, it is critical to distinguish between (i) the dropout genes where transcripts are missed in sequencing; and (ii) the “structural zeros” where the genes are truly un-expressed. Modeling the dropout events is especially challenging because of their complicated dependency on gene expression levels and cell characteristics. Specifically, dropouts are more likely to occur in genes expressed at low levels, and certain cells may have systematically higher dropout probabilities than others. In addition to dropout events, other challenges in modeling single cell data include the over-dispersion due to both cellular and technical variation, as well as high magnitude outliers due to bursts and fluctuations of gene expression levels. We refer the readers to Haque et al. (2017) for a more comprehensive review.

Despite the success of many early single-cell studies, statistical tools that account for the technical noise in single cell RNA-seq data, especially the dropout events, are limited. There have been efforts to analyze single cell data for various purposes. Many methods propose to quantify and account for technical noise using spike-ins (Brennecke et al., 2013; Vallejos, Marioni and Richardson, 2015; Vallejos, Richardson and Marioni, 2016). However, spike-ins are usually unavailable in single cell data due to its expenses in practice. For differential expression analysis, SCDE (Kharchenko, Silber-stein and Scadden, 2014) is based on a Bayesian hypothesis testing procedure using a three-component mixture model to capture technical noise; subsequently, MAST (Finak et al., 2015) uses a hurdle model that can adjust for various covariates; more recently, Vu et al. (2016) construct a beta-poisson mixture model, integrated within a generalized linear model framework. Various relevant problems have also been studied, including inferring the spatial localization of single cells in complex tissues (Satija et al., 2015), dimension reduction using Zero-Inflated Factor Analysis (ZIFA) (Pierson and Yau, 2015), and clustering unlabeled single cells while accounting for technical variation (Prabhakaran, Azizi and Peer, 2016). All of these aforementioned methods have been successfully applied to different single cell data sets. However, analytical methods that aim at the fundamental problem of imputing dropout genes and estimating the cell-type-specific gene expression profiles remain underdeveloped.

In this paper, we propose to jointly analyze single cell and bulk RNA-seq data using the Unified RNA-Sequencing Model (URSM), which simultaneously corrects for the dropout events in single cell data and performs deconvolution in bulk data. We point out that URSM only requires consistent cell types between both data sources, preferably measured on the same tissue from subjects with similar ages. It does not require the single cell and bulk data being measured on the same subjects, nor does it assume the same proportions of cell types in both data sets. Given a single cell data set, usually there are existing bulk data measured on the same tissue that can be modeled jointly using URSM. For example, BrainSpan provides extensive gene expression data on adult and developing human brains (Sunkin et al., 2013), and GTex establishes a human RNA-seq gene expression database across 43 tissues (GTEx Consortium, 2013).

By integrating single cell and bulk RNA-seq data, URSM borrows the strength from both data sources, and is able to (i) obtain reliable estimation of cell type specific gene expression profiles; (ii) infer the dropout entries in single cell data; and (iii) infer the mixing proportions of different cell types in bulk samples. Our framework explicitly models the dropout events in single cell data, and captures the relationship between dropout probability and expected gene expression levels. By involving high-quality bulk data, URSM achieves more accurate estimation of cellular expression profiles than using only single cell data. By incorporating the single cell data, URSM provides, for the first time, deconvolution of the bulk samples without going through the error-prone procedure of estimating marker genes. To the best of our knowledge, this is the first model that jointly analyzes these two types of RNA-seq data. We will illustrate in simulation (Section 4) and real-world data (Section 5) that URSM successfully corrects for the dropouts in single cell data, and provides reliable deconvolution for bulk samples.

## 2. A Unified Statistical Model

Suppose RNA-sequencing is conducted on *N* genes and *K* types of cells are of interest. Then bulk and single cell RNA-seq data can be linked together by a common profile matrix *A* ∈ ℝ^*N*×*K*^, where the *k*-th column *A*._*k*_ represents the expected *relative* expression levels of *N* genes in the *k*-th type of cells, such that each column sums to one. Note that by considering the *relative* expression levels, the profile matrix *A* does not depend on sequencing depths, and thus remains the same in both data sources. The two data sources provide two different views on the profile matrix *A*. In single cell data, the observations are independent realizations of different columns of *A* with extra noise due to dropout events. In bulk data, the expected relative expression levels for a mixture sample are weighted sums of columns of *A*, where the weights correspond to mixing proportions of different cell types. Here, we propose URSM to analyze the bulk and single cell RNA-seq data together, which borrows the strength from both data sets and achieves more accurate estimation on the profile matrix. This further enhances the performance of deconvolving bulk samples, as well as inferring and imputing the dropout genes in single cells.

The plate model of URSM for generating single cell and bulk RNA-seq data is given in Figure 1. Specifically, for single cell data, let *Y* ∈ ℝ^*N×L*^ represent the measured expression levels of *N* genes in *L* single cells, where the entries are RNA-seq counts. To model the dropout events, we introduce the binary observability variable *S* ∈ {0,1}^*N×L*^, where *S_il_* = 0 if gene *i* in cell *l* is dropped out, and *S_il_* = 1 if it is properly amplified. For each cell *l*, let *G_l_* ∈ {1, … ,*K*} denote its type, then the vector of gene expression *Y._l_* ∈ ℝ^*N*^ is assumed to follow a Multinomial distribution with probability vector *p_l_*, and the sequencing depth 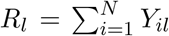 is the number of trials. Without dropout events, *p_l_* would be the corresponding column of the profile matrix, 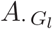, which is the true relative expression levels for cell type *G_l_*. With the existence of dropouts, *p_l_* becomes the element-wise product of 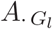 and *S._l_*, which is then normalized to sum to one. To capture the dependency between dropout probabilities and gene expression levels, the observation probability 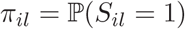 is modeled as a logistic function of 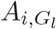,

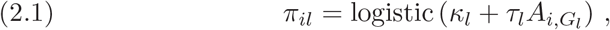

so that lowly expressed genes have high probabilities of being dropped out, where the coefficients (*κ_l_*, *τ_l_*) are cell-dependent that capture the cellular heterogeneity. Under this model, the set of dropout entries and structural zeros are defined as

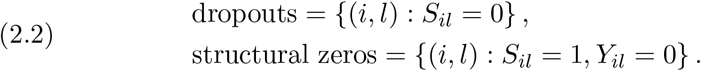

For bulk data, let *X* ∈ ℝ^*N*×*M*^ represent the RNA-seq counts of *N* genes in *M* bulk samples. For the *j*-th bulk sample, let *W._j_* ∈ ℝ^*K*^ denote the mixing proportions of *K* cell types in the sample, satisfying 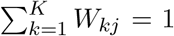. Then the gene expression vector *X._j_* ∈ ℝ^*N*^ is assumed to also follow a Multinomial distribution, where the probability vector is the weighted sum of *K* columns of *A* with the weights being *W._j_*, and the number of trials is the sequencing depth for sample *j*, defined as 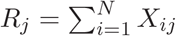.

For the hierarchical model setting, we assign the conjugate Dirichlet prior for the mixing proportions *W._j_*, and Gaussian priors for the cell-dependent dropout parameters (*κ_l_*, *τ_l_*). Here, we adopt an empirical Bayes approach, where the parameters are estimated by maximum-likelihood-estimations (MLE) using the expectation-maximization (EM) algorithm. Using this framework, our goal is threefold: (i) learn the profile matrix *A* as part of the model parameters, which characterizes the cellular gene expression profiles; (ii) make posterior inference on the dropout status *S* for single cell data, which can be used to identify dropout entries, and (iii) make posterior inference on the mixing proportions *W* in bulk samples. Finally, the inferred dropout entries in single cell data can be imputed by their expected values using the estimated *A* and sequencing depths *R_l_*.

**Fig. 1.**
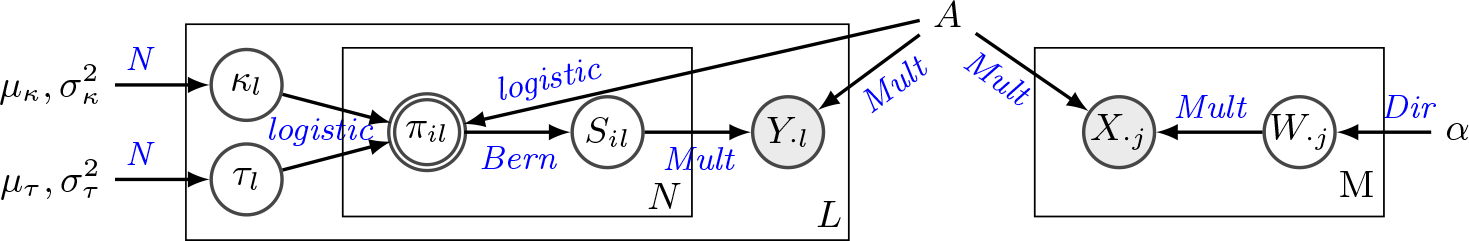
*Plate model of URSM, with both single cell data (on the left) and bulk samples (on the right). The two greyed nodes X and Y represent observed gene expression levels. Node S is a binary variable representing dropout status in single cells, and node W represents the mixing proportions in bulk samples. The node π representing observation probability is double-circled because it is deterministic, and all model parameters are shown without circles, including the profile matrix A that links the two data sources.*

*Full model specification‥*

- Bulk data
  − 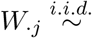 Dirichlet(*α*) for *j* = 1, … , *M*, where *α* ∈ ℝ^*K*^, *α* ≥ 0.
  − 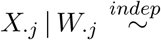 Multinomial(*R_j_*, *AW._j_*) for *j* = 1, … , *M*, where 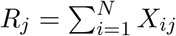
- Single cell data
  − 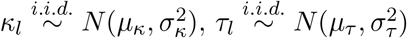 for *l* = 1, … , *L*.
  − *π_il_* = logistic 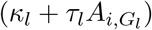, where *G_l_* ∈ {1, … , *K*} is the type of the *l*-th cell.
  − 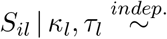 Bernoulli(*π_il_*) for *i* = 1, … , *N*; *l* = 1, … , *L*.
  − 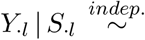 Multinomial(*R_l_*, *p_l_)* for *l* = 1, … , *L*, where 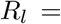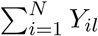

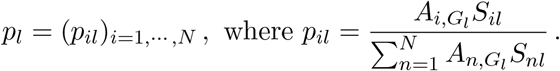

### Remark 1.

We assume all entries in *A* to be strictly positive. In principle, one can allow some entries *A_ik_* to be exactly zero, but this will lead to a degenerate multinomial distribution and complicate the likelihood function. In addition, making inference on *S_il_* when 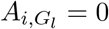 is an ill-defined problem. If *A_ik_* = 0, then we will have *X_il_* = 0 for all type-*k* cells, but such structure rarely appears in real data. In practice, it is usually helpful to use some small positive numbers rather than exact zeros to capture the background signal in sequencing processes (Kharchenko, Silberstein and Scadden, 2014).

### Remark 2.

It is straightforward to use one part of URSM when only one data source is available. In Section 4, we will show the performance of the submodel for single cell data. It is also possible to use the submodel for bulk data when only bulk data are available, but extra information about marker genes needs to be incorporated in this scenario to avoid the non-identifiability issue, as explained in Section 1.

## 3. Inference and Estimation: EM Algorithm

This section presents an expectation-maximization (EM) algorithm (Dempster, Laird and Rubin, 1977) for fitting the maximum likelihood estimation (MLE) of the parameters *θ* = (*A*,*α*,*μ_κ_*,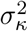,*μ_τ_*, 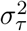), as well as a Gibbs sampling algorithm for posterior inference on latent variables *H* = (*W, S*, *κ*, *τ*). As illustrated in Section 2, the key values of scientific interests include (i) an estimate of the profile matrix *A* that characterizes the cellular gene expression profiles; (ii) 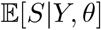, the inferred dropout probability at each entry in single cell data; and (iii) 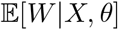, the inferred mixture proportion of bulk samples.

The main difficulty of handling our model is the intractable posterior distributions due to non-conjugacy. Therefore, approximate inference needs to be performed. One of the main methods for approximate inference in Bayesian modeling is Monte Carlo Markov Chain (MCMC) sampling (Gelfand and Smith, 1990), where a Markov chain on latent variables is constructed, with stationary distribution being the true posterior. After obtaining a long enough chain, the posterior can be approximated with empirical estimation. Gibbs sampling (Geman and Geman, 1984; Casella and George, 1992) is one of the most widely used forms of MCMC algorithms given its simplicity and efficiency. On the other hand, variational methods form an alternative line for approximate inference, where the posterior is approximated analytically by a family of tractable distributions (Jordan et al., 1999; Wainwright and Jordan, 2008; Blei, Kucukelbir and McAuliffe, 2016). While being computationally scalable in many large-scale problems, variational methods are inherently less accurate due to the inevitable gap between the variational distributions and the true posterior distribution.

In this paper, we present a Gibbs sampling algorithm for approximate inference on latent variables using the data augmentation trick. This algorithm can also be used in the E-step of the EM procedure, leading to a Gibbs-EM (GEM) algorithm for obtaining MLEs of model parameters (Dupuy and Bach, 2016). The specific steps are outlined in Section 3.1 and Section 3.2, and more details can be found in the supplement. Finally, we point out that one can also proceed with variational inference, but due to space limitation, we do not pursue this approach in detail.

### 3.1. *E-step: Gibbs sampling.*

The latent variables for bulk data and single cell data are conditionally independent given observed data *X, Y* and parameters. Therefore, Gibbs sampling can be performed on the two data sources in parallel. In this section, we describe the sampling procedure for the two parts separately.

*Bulk data.* To obtain the posterior inference of *W* (the mixing proportions) in bulk data, we re-write the model to be mixture of multinomials by introducing the augmented latent variables *Z* and *d* as follows:

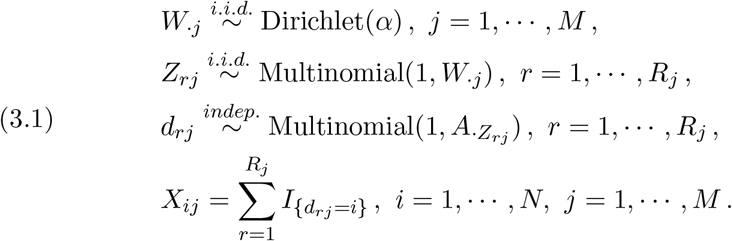

Note that this model is closely related to the Latent Dirichlet Allocation (LDA) model (Blei, Ng and Jordan, 2003) in topic modeling, if we view a gene as a word, a cell type as a topic, and a bulk sample as a document. Although the Gibbs sampling algorithm has been developed for LDA in Griffiths and Steyvers (2004), there are two difficulties that prevent us from directly applying this algorithm to our model. First, the LDA model assumes observations of *d_rj_*, which are the actual words in an document, but in RNA-seq data, only the final counts *X_ij_* are observed. Second, the sequencing depths *R_j_*’s are typically large in real data, so it will be extremely computationally demanding to keep track of *Z_rj_* and *d_rj_*. Therefore, we propose a modified algorithm by defining another set of augmented latent variables

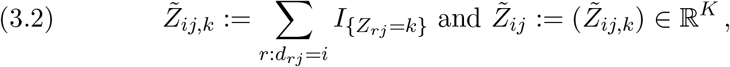

and it can be shown that

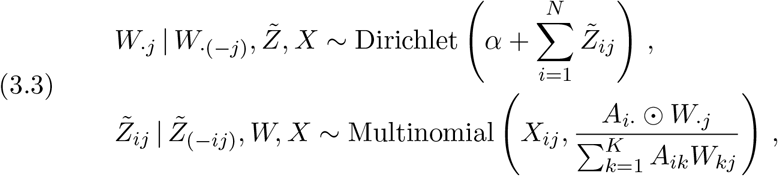

where ⨀ denotes element-wise multiplication, and the index (−*i*) denotes everything else other than *i*.

*Single cell data.* As for posterior inference of *S*, *κ*, *τ* in single cell data, note that the first part of the model can be re-written as

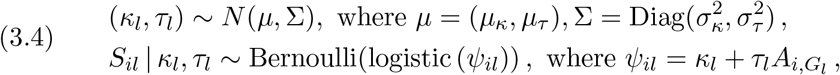

which has the same form as a Bayesian logistic regression, with covariates being 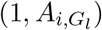. Therefore, following the recent development of Gibbs sampling technique in this area (Polson, Scott and Windle, 2013), we introduce a set of augmented latent variables *ω*, and the conditional complete posteriors can be shown to be

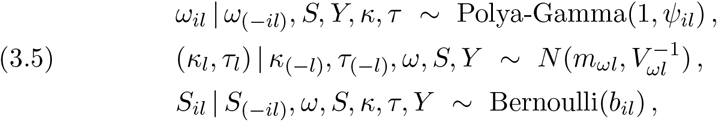

where

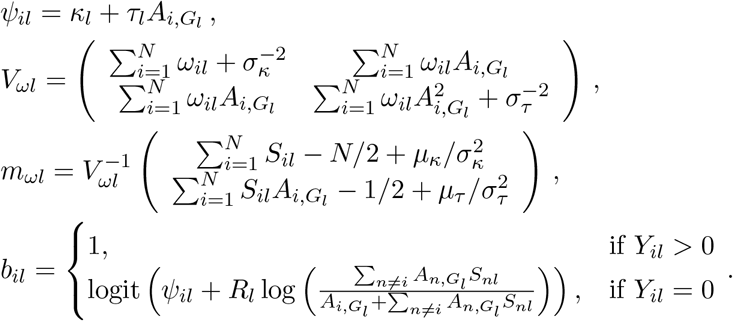

### 3.2. *M-step.*

In the M-step of GEM algorithm, the parameters are updated to maximize a lower bound on the expected complete log likelihood function, or the so-called Evidence Lower BOund (ELBO), where the posterior expectation 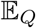 is estimated using Gibbs samples obtained in the E-step.

The optimal dropout parameters (*μ_κ_*,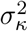,*μ_τ_*, 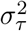) have the following closed forms:

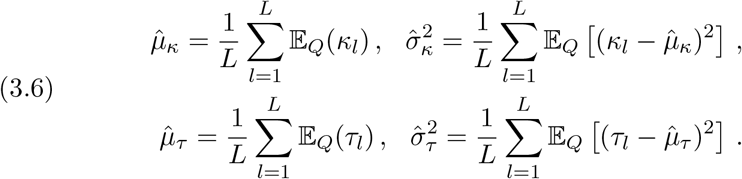

For *A* and *α*, there are no closed form solutions, and we use the projected gradient ascent algorithm:

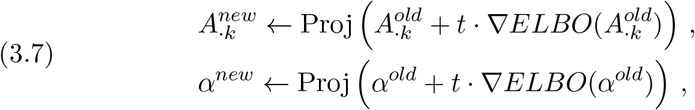

where the step size *t* is determined by backtracking line search, and the *Proj* function is the projection onto the feasible set:

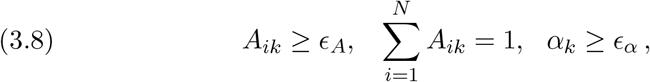

where ϵ_*A*_, ϵ_*α*_ > 0 are some small pre-determined constants. The gradients are computed as

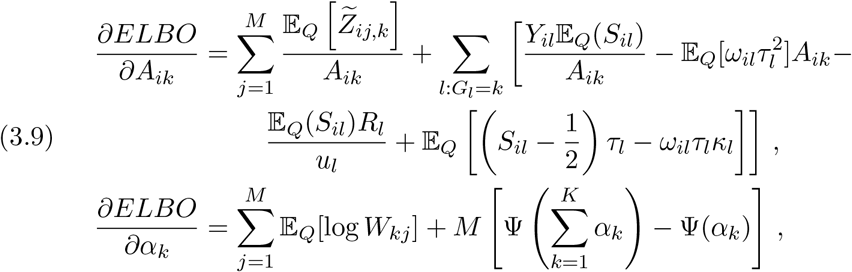

where 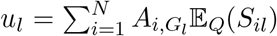 and ψ(·) is the digamma function. More detailed derivations can be found in the supplement.

## 4. Simulation Results

In this section, we evaluate the performance of URSM in synthetic datasets. We focus on the accuracy of recovering the profile matrix *A* and mixing proportions *W*, as well as the ability of distinguishing between dropout entries and structural zeros using the posterior inference of *S*.

### 4.1. *Settings.*

Let *N* be the number of genes. The sequencing depths for bulk samples are independently generated from Poisson(50 *N*). To account for the fact that the sequencing depths of single cell data are usually much lower and highly variable, they are generated from Negative Binomial with mean 2 *N* and dispersion parameter 2.

The cell-type specific profile matrix *A* is generated as follows: (i) simulate all entries independently from log-normal with *μ* = 0, *σ* = 1; (ii) for each cell type *k*, let *N_m_* = 10 genes be marker genes, i.e., set *A_il_* = 0 for *l* ≠ *k*; (iii) for each cell type *k*, let *N_a_* = 10 genes be anti-marker genes, i.e., set *A_ik_* = 0; (iv) let another set of *N_h_* = 30 genes be house-keeping genes that have same expression levels in all cell types; (v) finally, properly normalize A so that each column sums to 1. Specifically, in each column, we normalize the *N_h_* house-keeping genes such that they sum to *N_h_/N*, and the remaining genes sum to 1 − *N_h_/N*.

Finally, the observation status {*S_il_*}_*il*_ for each gene *i* in each single cell *l* is simulated independently from Bernoulli(*π_il_*). Recall that *S_il_* = 0 indicates a dropout, and the dropout probability is determined by

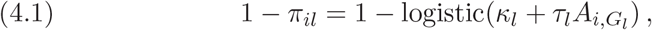

where *G_l_* ∈ {1,…, *K*} is the type of cell *l*. In the following sections, *κ_l_*’s are independently generated from Normal(−1,0.5^2^), and *τ_l_*’s are independently generated from Normal(1.5 *N*, (0.15 *N*)^2^). Note that by construction, the mean of each column of *A*, *Ā*.,_*k*_, is always 1/*N*. Therefore, 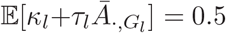 for each cell, which corresponds to an average dropout probability of 37.8%. and the maximal dropout probability is 73.1% when *A_ik_* = 0.

### 4.2. *Estimation of profile matrix.*

In this section, we illustrate that URSM provides accurate estimation on the profile matrix *A* after correcting for dropouts and utilizing bulk samples. Following the simulation settings specified in Section 4.1, we generate *L* = 100 single cells and *M* = 150 bulk samples on *N* = 200 genes. We consider *K* = 3 cell types. For single cells, 30%, 30% and 40% of the cells are assigned to the 3 different types, respectively. For bulk samples, the hyper parameter of the mixing proportions is set to *α* = (1, 2, 3). The dropout probability curves, simulated following equation (4.1), are shown in Figure 2a. The simulated single cell data has 64.6% entries being zero.

**Fig. 2.**
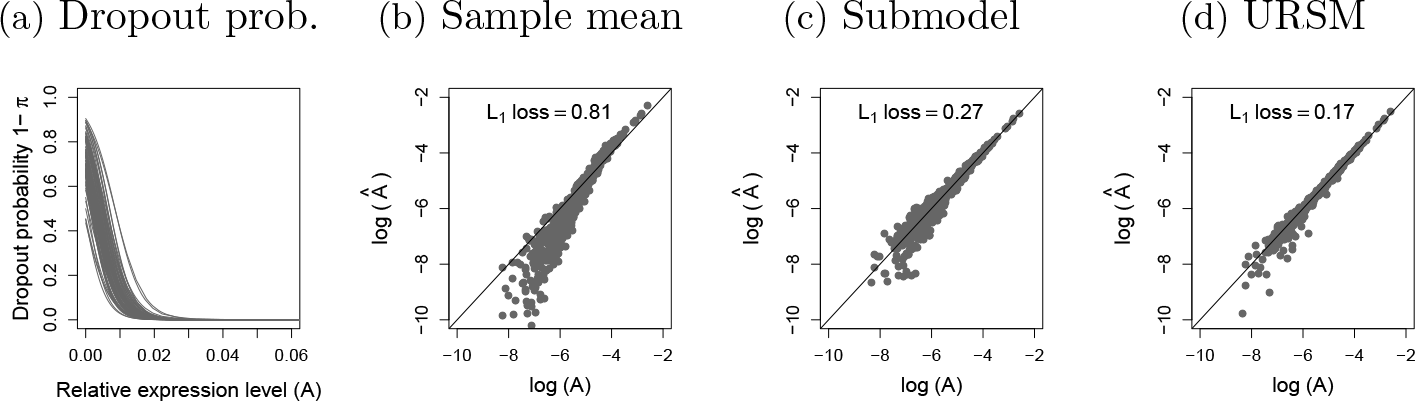
**(a)** *Simulated logistic dropout probability curves for 100 single cells, as defined in equation (4.1).* **(b)** − **(d)** *True profile matrix **A** versus the estimated **Â**, plotted in the log scale, using (i) the naive sample mean estimation (equation (B.6)); (ii) a submodel using only single cell data; (iii) URSM with both single cell and bulk data. The *L*_1_ loss* Σ_*i,k*_ | *Â_ik_ − A_ik_* | *is reported on the top.*

A naive method to estimate the profile matrix *A* is to use the sample means of single cell expression levels, after normalizing by their sequencing depths. Specifically, recall that *Y* ∈ ℝ^*N*×*L*^ represents the observed expression levels in single cells, {*G_l_*}_*l*=1,…,*L*_ represent the cell types, and {*R_l_*}_*l*=1,…,*L*_ are the sequencing depths, defined as *R_l_* = Σ_*i*_ *Y_il_*. Then an entry *A_ik_* can be estimated by

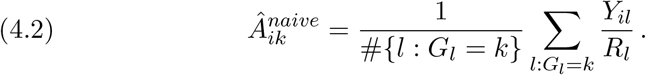

However, due to the presence of dropout events and the dependency between *π_il_* and *A*, this naive sample mean estimation is biased, with *L*_1_ loss 0.81 (Figure 2b), where the *L*_1_ loss is computed as Σ_*i,k*_ | *Â_ik_ − A_ik_* |. On the other hand, by explicitly modeling the occurrence of dropout events and capturing the relationship between dropout probability and expected expression level, a submodel of URSM that only uses single cell data successfully corrects for the bias, and substantially reduces the loss to 0.27 (Figure 2c). Finally, by integrating the bulk data, URSM further improves the estimation and further reduces the *L*_1_ loss to 0.17 (Figure 2d).

### 4.3. *Deconvolution of bulk samples.*

Now we further examine the model performance on inferring the mixing proportions W in bulk samples, using the same simulation setting as in Section 4.2. We compare the performance of URSM to three widely used deconvolution methods: Digital Sorting Algorithm (DSA) (Zhong et al., 2013), semi-supervised Nonnegative Matrix Factorization (ssNMF) (Gaujoux and Seoighe, 2012), and Cibersort (Newman et al., 2015).

Both DSA and ssNMF rely heavily on a set of *given* marker genes as input to guide the matrix factorization, where a “marker gene” is only expected to express in one cell type. Unfortunately, marker genes are rarely known in practice, and a widely adopted procedure is to estimate the list of marker genes from purified cells by selecting those with the most different expression levels across cell types. Here, we mimic this procedure by estimating a list of marker genes from single cell data to guide DSA and ssNMF. Specifically, we adopt the method in Abbas et al. (2009), which calculates a *p*-value of each gene by comparing its expression level in the highest and second-highest types of cells, then selects the group of genes with the smallest *p*-values. Figure 3 shows the *L*_1_ loss of estimating *A* and *W* using DSA and ssNMF with different sets of estimated marker genes with *p*-values smaller than {10^−8^, … , 10^−3^}, and the number of selected marker genes is listed in Table 1. It is clear that these two algorithms are sensitive to the input marker genes. For comparison, we also evaluate the performances of DSA and ssNMF when the oracle information of true marker genes is available.

On the other hand, Cibersort requires a “signature” matrix containing the expression levels of a group of “barcode” genes that collectively distinguish between different cell types. Note that this essentially requires knowing part of the profile matrix *A*, which contains much more information than the marker gene list. Here, we use the estimated *Â* from our unified model as the signature matrix for Cibersort. We report the *L*_1_ loss of estimating *W* when Cibersort only takes the expression levels of the selected marker genes, as well as when Cibersort uses the entire *Â*. Figure 3b suggests that Cibersort prefers larger number of barcode genes as input.

Finally, URSM automatically utilizes the information in single cell data to guide deconvolution. Figure 3 illustrates that URSM and Cibersort usually outperform DSA and ssNMF using estimated marker genes, and achieve comparable *L*_1_ loss even when DSA and ssNMF have the oracle information of marker genes.

**TABLE 1.** *Number of selected, marker genes using different thresholding p-values.*

| # of markers | $\log_{10}(\text{p-value})$ | | | | | | True markers |
| --- | --- | --- | --- | --- | --- | --- | --- |
|  | -8 | -7 | -6 | -5 | -4 | -3 |  |
| cell type 1 | 5 | 8 | 11 | 16 | 19 | 27 | 10 |
| cell type 2 | 2 | 2 | 8 | 11 | 16 | 23 | 10 |
| cell type 3 | 1 | 2 | 7 | 10 | 17 | 21 | 10 |

### 4.4. *Inference of dropout entries in single cell data.*

Next, we present the inference on dropout entries in single cell data, again using the same setting as in Section 4.2. Here, our goal is to distinguish between dropout entries and structural zeros, as defined in equation (2.2). Note that we only need to make inference for locations where the observed expression levels are zero, i. e., on the set {(*i, l*) : *Y_il_* = 0}. Recall that *S_il_* = 0 if gene *i* is dropped out in cell *l*, and our model provides the estimated posterior mean of *S*:

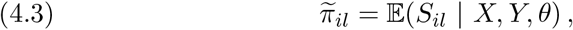

where *θ* denotes the model parameters. Hence a natural approach is to predict the entries with small 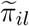 to be dropouts.

**Fig. 3.**
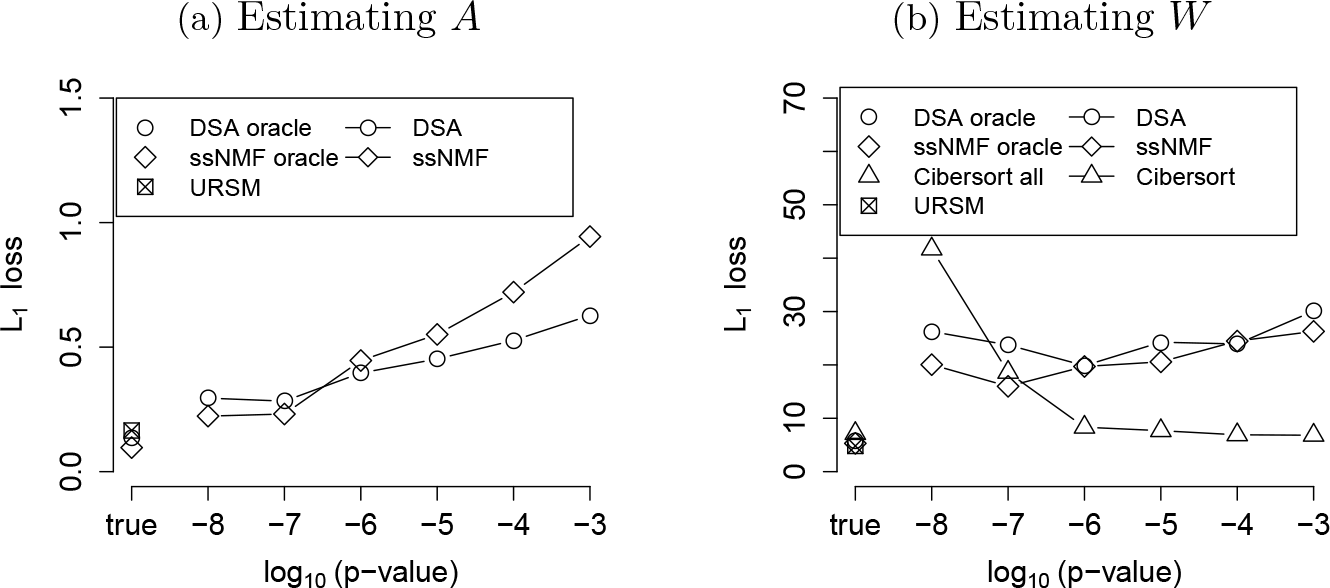
*The L_i_ loss of recovering* **(a)** *the profile matrix*, Σ_*i,k*_ | *Â_ik_ − A_ik_* |, *and* **(b)** *mixing proportions*, Σ_*k,j*_ | *Ŵ_kj_ − W_kj_* |. *We evaluate DSA and ssNMF when the marker genes are extracted from single cell data using different thresholds of p-values, as well as under the oracle condition where the true marker genes are given. We evaluate Cibersort on estimating W when the input signature matrix is based on the estimated Â from URSM. We report its performance when the entire Â is used (“Cibersort all”), as well as when only the estimated marker genes are used (“Cibersort”). The performance of URSM is plotted with a square in both panels, which does not depend on thresholding p-values.*

A potential competitor for imputing dropout entries is the Nonnegative Matrix Factorization (NMF) (Lee and Seung, 2001). One can construct a low-rank approximation to the single cell expression matrix *Y* ∈ ℝ^*N*×*L*^ using NMF. Intuitively, the approximated values tend to be higher at dropout entries, and closer to zero at structural-zero entries. As shown in Figure 4a, if the rank is properly specified, this simple NMF-based method demonstrates certain ability to distinguish between dropout genes and structural zeros, but not as well as URSM. In addition, in order to further impute the dropout entries, a good estimation of the profile matrix *A* is also needed. Figure 4b shows the estimation of *A* by taking sample average as in equation (B.6), with *Y* substituted by the NMF approximation. It is clear that the NMF approach fails to correct for the bias introduced by the dropout events, while URSM succeeds in both identifying dropout entries and obtaining an unbiased estimation of *A* (recall Figure 2d).

**Fig. 4.**
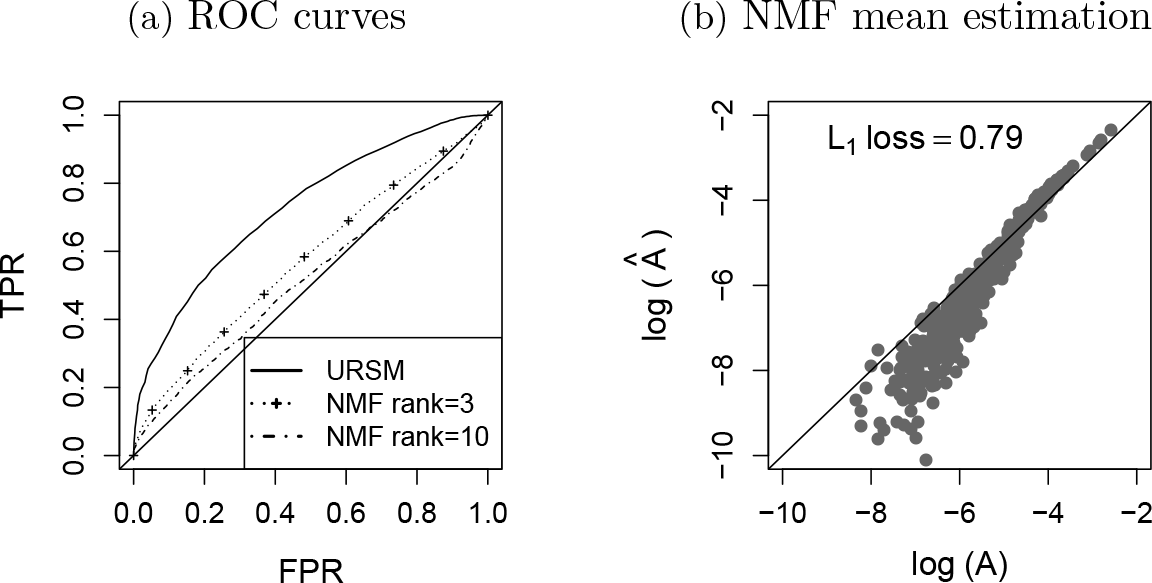
**(a)** *ROC curves of identifying dropout entries in single cell data.* **(b)** *True profile matrix A versus the sample average of a rank-3 NMF approximation, plotted in the log scale. The L_i_ loss* Σ_*i,k*_ | *Â_ik_ − A_ik_* | *is reported on the top.*

### 4.5. *Robustness.*

Finally, we demonstrate the robustness of our model. We apply URSM under the scenario where the number of cell types in single cell data *K_sc_* is not equal to the number of cell types in bulk data *K*_bk_, as well as when the number of genes *N* grows. URSM always takes *K* = max{*K_sc_*, *K_bk_*} as input, and estimates *Â_unif_* ∈ ℝ^*N*×*K*^ and *Ŵ_unif_* ∈ ℝ^*K*×*M*^. When *K_sc_* > *K_bk_*, it is straightforward to directly apply URSM, and ideally the estimated *Ŵ_unif_* will assign zero proportions to the missing cell types in bulk samples. However, when *K*_*sc*_ < *K_bk_*, without extra information, deconvolution is an ill-defined problem because of the non-identifiability issue (see Section 1 for more details). In order to find a biological meaningful solution, we follow the idea in ssNMF (Gaujoux and Seoighe, 2012) and use a set of marker genes to initialize the parameters for the EM algorithm by setting the corresponding entries in *A* to be zero. We consider the scenario where for each cell type, 5 true marker genes and 3 imperfect marker genes are used for initialization. The imperfect marker genes are selected from the non-marker genes, where we pick the ones with the largest difference between the highest and second highest expression levels across cell types in *A*.

Following Section 4.1, we simulate *M* = 150 bulk samples, where the mixing proportions in bulk samples are generated from Dir(*α*) with *α* = (1,…,*K_bk_*). For single cell data, we generate 40 cells in the majority cell type, and 30 cells in each of the remaining *K_sc_* − 1 types. To reduce the computation load and enhance stability, we use the maximum a posteriori estimation for *W* in the E-step for bulk samples. More details are included in the supplement.

Again, we compare URSM to DSA, ssNMF, and Cibersort. Both DSA and ssNMF require a set of marker genes as input, and we report their performances under two scenarios: (i) the oracle scenario where 5 true marker genes are provided for each cell type; and (ii) a more realistic scenario as used by our uniform model, where 5 true marker genes and 3 imperfect marker genes are provided for each cell type. Note that when *K_sc_* > *K_bk_*, bulk samples contain no information of the expression patterns for the missing cell types, so we allow DSA and ssNMF to only deconvolve *K_bk_* cell types in these cases. We point out that this strategy favors the DSA and ssNMF methods by providing them extra information of the missing cell types in bulk samples. For Cibersort, as in the previous sections, we use the estimated profile matrix obtained from our uniform model as the input signature matrix.

Figure 5a summarizes the performance of different models under various choices of *K_sc_* and *K_bk_* when *N* = 200 in 10 repetitions. In order to make a comparable comparison across different *K*’s, we report the *average per cell type L*_1_ loss, i.e., the average *L_1_* loss ‖*Â*.,_*k*_ − *A*.,_*k*_‖_1_ and ‖*Ŵ*.,_*k*_ — *W*,_*k*_‖_1_ across all columns *k*. We see that the performance of URSM remains robust under different settings, and is usually comparable to DSA and ssNMF algorithms even when the latter two algorithms have the oracle marker gene information. Not surprisingly, Cibersort has similar performance as URSM because it uses our estimated *Â_unif_* as input. We point out that when the sample mean estimation *Â_naive_* (equation (B.6)) is given to Cibersort as the signature matrix, the performance is unstable and it cannot provide deconvolution when *K_sc_* < *K_bk_*. Finally, we also demonstrate the performance of different models when *N* = {200, 500,1000}, where we set *K_sc_* = *K_bk_* = 3. Figure 5b verifies that URSM remains robust with larger numbers of genes.

## 5. Application to Fetal Brain Data

### 5.1. *Data pre-processing.*

In this section, we apply URSM to gene expression measured on fetal brains. The single cell RNA-seq data comes from Camp et al. (2015), where 226 cells from fetal brains are sequenced on 18,927 genes. The authors have removed endothelial cells and interneurons, and the remaining 220 single cells are labeled into three types: 40 apical progenitors (APs), 19 basal progenitors (BPs), and 161 neurons (Ns). In addition, the authors have normalized the RNA-seq counts using FPKM (Fragments Per Kilobase of exon per Million fragments mapped) and performed log-transformation by log_2_(*x* + 1). We refer the readers to Camp et al. (2015) for more details of the single cell data pre-processing. On the other hand, microarray bulk gene expression data on fetal brains is provided by the BrainSpan atlas (Kang et al., 2011). Within the same window of development, 12 to 13 post-conception week, 72 bulk samples from prefrontal cortex are measured on 16,947 genes. To apply our model, the single cell RNA-seq data are transformed back to linear scale by 2^*x*^ − 1, and all measurements are truncated to integers. To approximate the RNA-seq counts in bulk samples, we transform the BrainSpan microarray data in the same way and treat them as pseudo-RNA-seq counts. The resulting bulk samples have an average pseudo sequencing depth of 5.5 × 10^6^, which is 26 times larger than the average effective sequencing depth in single cells, 2.1 × 10^5^, where the effective sequencing depth is calculated as the sum of FPKM across all genes in each single cell.

**Fig. 5.**
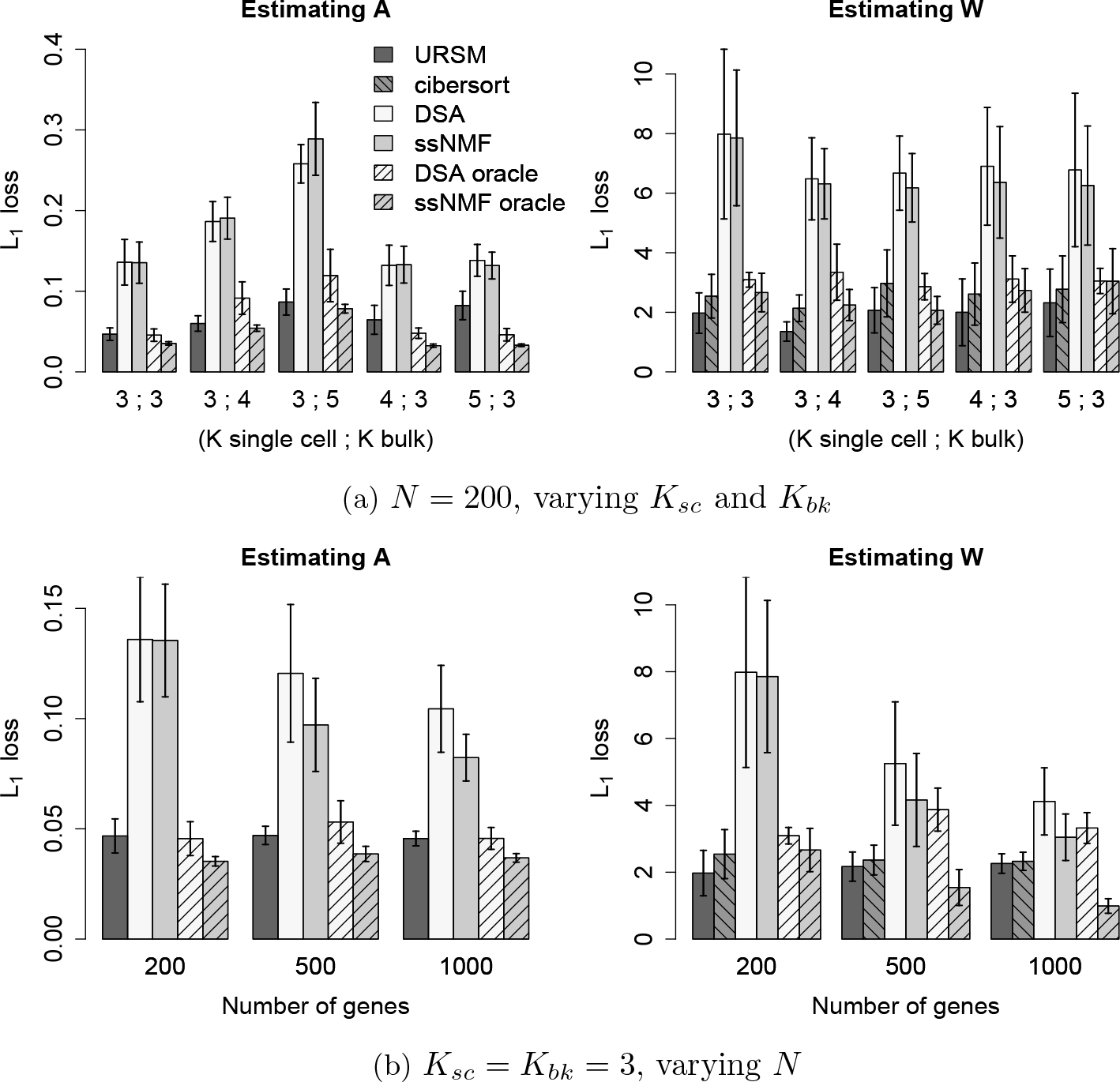
*The average per cell type L_1_ loss of recovering the profile matrix A and the mixing proportions W in 10 repetitions, with the standard deviations shown by the error bars, when* **(a)** *K_sc_,K_bk_* ∈ {3,4, 5} *with N* = 200 *genes;* **(b)** *N* = {200, 500,1000} *with K_sc_* = *K_bk_* = 3. *Each figure shows the performance of (i) URSM; (ii) DSA and ssNMF with 5 true marker genes and 3 imperfect marker genes per cell type as input; (iii) DSA and ssNMF under the oracle scenario where 5 true marker genes per cell type are provided. We also report the performance of Cibersort for estimating W using the estimated Â_unif_ from URSM as the input signature matrix.*

To reduce computational load, we only focus on genes with significantly different expression levels among the three cell types. Specifically, we use the 315 so-called PC genes proposed in Camp et al. (2015), which have the largest loadings in a Principal Component Analysis (PCA) and account for the majority of cellular variation. After restricting to the overlapping genes that are also sequenced in BrainSpan bulk samples, a final list of 273 PC genes are obtained and used in the following analyses. When restricting to these 273 genes, the average effective sequencing depth (i.e., the sum of RNA-seq counts in each sample) is 3.2 × 10^5^(*sd* = 1.6 × 10^4^) in BrainSpan tissues, and 1.4 × 10^4^(*sd* = 4.3 × 10^3^) in single cells .

Due to the nature of active cell development from APs and BPs to Neurons in fetal brains, we expect to have a few cells that are actively transitioning between two cell types, whose labels are ambiguous. We first remove these ambiguously labeled cells from our analysis. Specifically, we project the single cells to the leading 2-dimensional principal subspace, where the pseudo developing time is constructed using the Monocle algorithm (Trapnell et al., 2014). Based on the results, the 3 BPs that are close to AP or Neuron clusters are removed, so are the 4 Neurons that are close to AP or BP clusters (Figure 6b). The remaining 213 single cells are retained for analysis, and their gene expression levels on the 273 PC genes are visualized in Figure 6a.

### 5.2. *Imputation of single cell data.*

Here, we apply URSM to identify and impute the dropout entries in single cell data. Note that in order to distinguish between dropout entries and structural zeros in single cell data (equation (2.2)), we only need to focus on the entries where the observed gene expression levels are zero. The inference of dropout entries is based on the estimated posterior expectation of 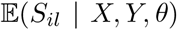. As a result, among the 37,771 zero-observation entries, 45.7% are inferred to be dropouts with probability one (Figure 6c). These entries are then imputed by their expected values, calculated using the corresponding entries in the estimated profile matrix *A* multiplied by the sequencing depths of the corresponding cells. To illustrate the impact of imputation, we apply PCA again on the imputed data. Figure 6d visualizes the cells in the first two principal components, and the clusters for different cell types are more clearly separated.

### 5.3. *Deconvolution of bulk samples.*

Finally, we present the deconvolution results of bulk samples using URSM. According to the prior knowledge that the proportions in bulk samples should be roughly consistent with that in single cell data, the mixing parameter *α* is initialized at (2 × 10^4^, 10^4^, 7 × 10^4^) for AP, BP and Neurons. The scale of *α* is chosen to be comparable to the average effective sequencing depths of 1.4 × 10^4^ among all single cells. Figure 7a shows the inferred mixing proportions of APs, BPs and Neurons in each of the 72 bulk samples, with an average of 17.7% AP cells, 8.7% BP cells and 73.6% Neurons.

For comparison, we also apply the Digital Sorting Algorithm (DSA) (Zhong et al., 2013), semi-supervised Nonnegative Matrix Factorization (ssNMF) (Gaujoux and Seoighe, 2012), and Cibersort (Newman et al., 2015) on the BrainSpan bulk samples. The marker genes for DSA and ssNMF are selected by comparing each gene’s expression level in the highest and second-highest types of cells in the single cell data, and genes with *p*-value < 10^−5^ are treated as markers (Abbas et al., 2009). This procedure leads to 21 AP markers, 6 BP markers and 28 Neuron markers, which serve as input to DSA and ssNMF. For Cibersort, the input signature matrix is provided by the estimated *Â* from URSM. Figures 7d to 7b suggest that the proportions estimated by ssNMF tend to have too large variations, while DSA overestimates the neural composition, and Cibersort obtains similar results as URSM.

As another perspective to verify the deconvolution results, we use the intuition that the true proportions of a cell type should be correlated with the expression levels of its marker genes in bulk samples. To check whether this holds in the results, we first normalize each bulk sample by their effective sequencing depths, such that the normalized expressions sum to one in each sample. We focus on 7 genes based on biological knowledge, including the radial glia (RG) markers *PAX6* and *GLI3* that are expected to only express in AP and BP cells, the RG marker *HES1* that is mostly expressed in AP cells, the early BP marker *HES6*, as well as neuronal genes *NEUROD6*, *BCL11B* and *MYT1L* (Camp et al., 2015). Table 2 summarizes the correlations calculated by estimated proportions using different methods, and we see that URSM and Cibersort usually achieve the highest correlations. Finally, we point out that if Cibersort uses the naive sample mean estimation from single cell data as the signature matrix, it will fail to identify BP cells and achieve much lower correlations.

**Fig. 6.**
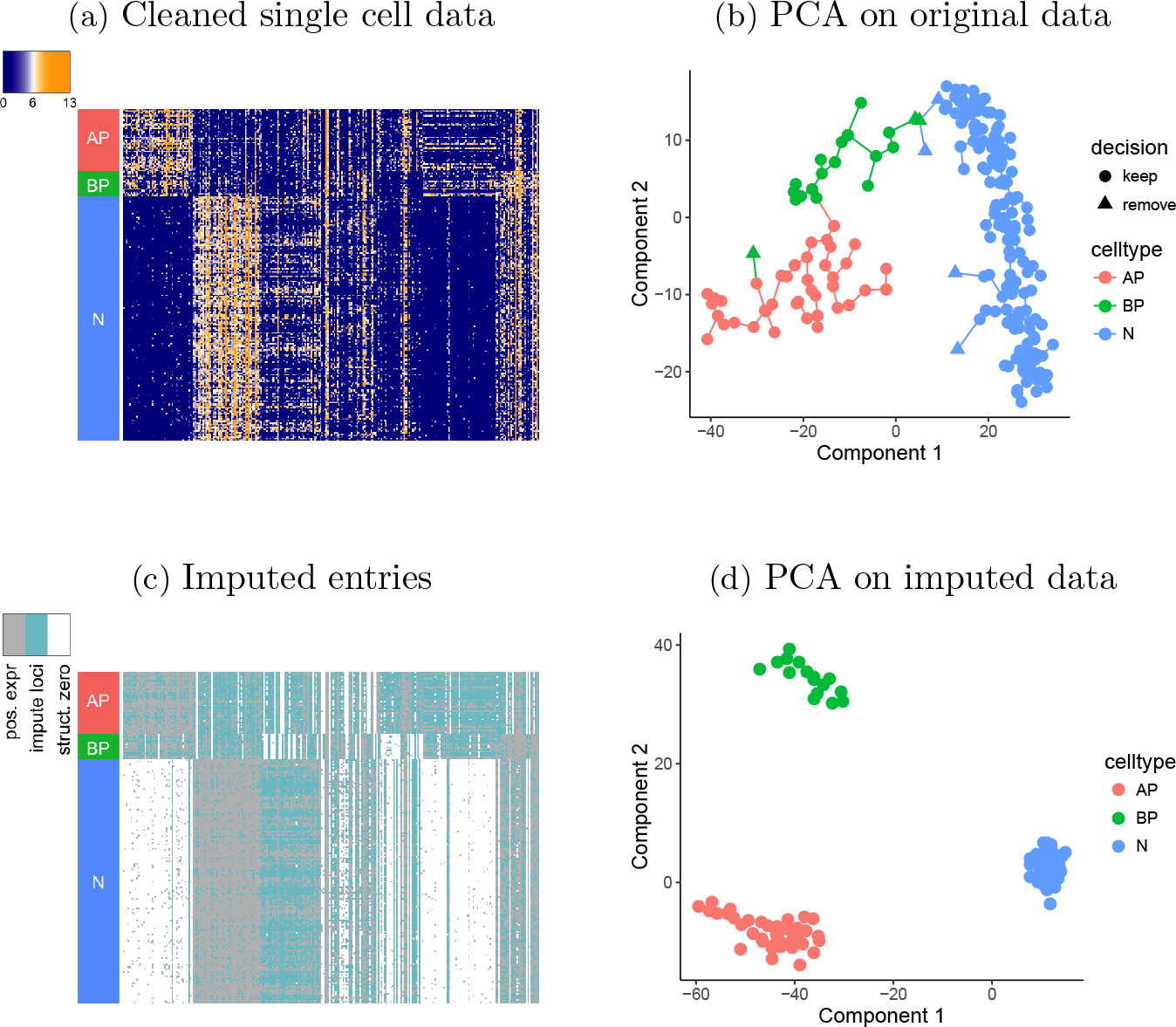
Fig. 6. **(a)** *Single cell gene expressions (*log_2_ *(FPKM+1)) after removing 7 ambiguously labeled cells. Rows are 213 cells and columns are 273 genes.* **(b)** *PCA applied on the original single cell data with 220 labeled cells using 273 PC genes, where the Monocle algorithm is applied to construct pseudo developmental times. 7 cells are identified to be ambiguously labeled and are removed from our analyses (marked as triangles).* **(c)** *Entries in cleaned single cell data that are inferred to be dropout and imputed (marked in blue) versus the entries that are inferred to be structural zeros (marked in white) in cleaned single cell data. The entries with positive expression levels have no need for posterior inference, and are marked in grey.* **(d)** *After imputing dropout genes, PCA is conducted on the 213 cells using 273 PC genes, and the three different types of cells are more clearly separated.*

**Fig. 7.**
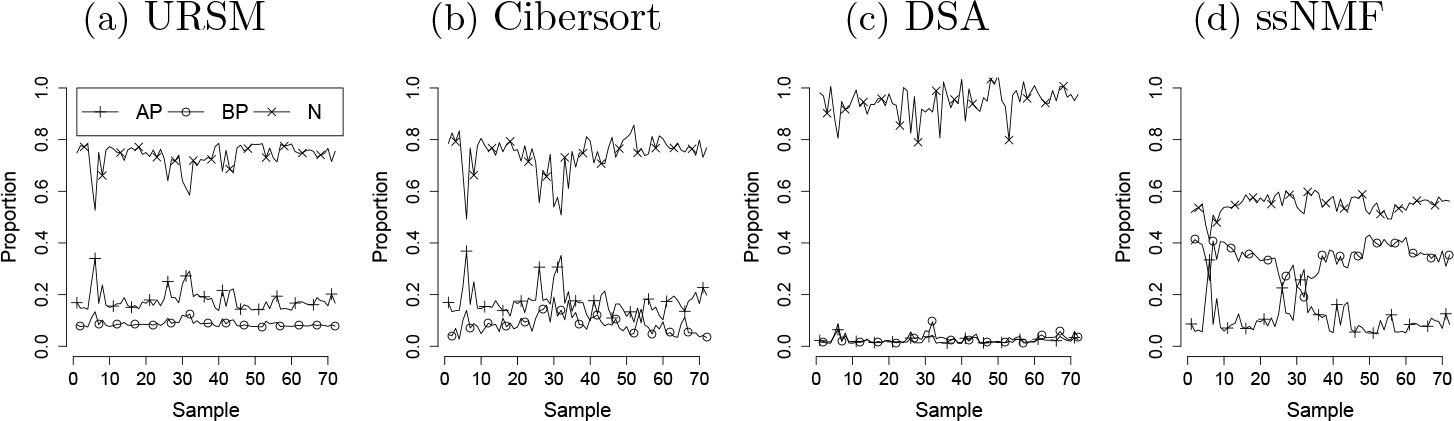
*Deconvolution of bulk samples into three cell types, using* **(a)** *URSM;* **(b)** *Cibersort;* **(c)** *Digital Sorting Algorithm (DSA);* **(d)** *semi-supervised Nonnegative Matrix Factorization (ssNMF).*

**TABLE 2.** *Correlation between the estimated proportions of a cell type k in bulk samples, (W_kj_)_j_, and the normalized expression levels (X_ij_/R_j_)_j_ of its marker gene i in bulk samples. For genes marking both AP and BP, the sum of proportions is used.*

| Gene | Marked cell type | URSM | Cibersort | DSA | ssNMF |
| --- | --- | --- | --- | --- | --- |
| HES1 | AP | 0.73 | 0.62 | **0.80** | 0.68 |
| HES6 | BP | **0.66** | 0.58 | 0.53 | -0.72 |
| PAX6 | AP.BP | **0.91** | 0.80 | 0.80 | 0.61 |
| GLI3 | AP.BP | **0.90** | 0.80 | 0.83 | 0.54 |
| NEUROD6 | N | 0.28 | **0.37** | 0.02 | -0.36 |
| BCL11B | N | 0.45 | **0.57** | 0.23 | 0.02 |
| MYT1L | N | **0.44** | 0.37 | 0.32 | 0.80 |

## 6. Discussion

In this paper, we propose URSM, a unified framework to jointly analyze two types of RNA-seq data: the single cell data and the bulk data. URSM utilizes the strengths from both data sources, provides a more accurate estimation of cell type specific gene expression profiles, and successfully corrects for the technical noise of dropout events in single cell data. As a side product, URSM also achieves deconvolution of bulk data by automatically incorporating the cellular gene expression patterns.

Dropouts present one of the biggest challenges to modeling scRNA-seq data. URSM assumes a dependency between expression level and the probability of observing dropout and aims, probabilistically, to infer which observations are likely dropouts. There are a number of alternative approaches in the literature; for a discussion see Huang et al. (2017b) and Vallejos et al. (2017). The most common statistical approach is to explicitly model the zero-inflation process, for example, SCDE (Kharchenko, Silberstein and Scadden, 2014), MAST (Finak et al., 2015) and ZIFA (Pierson and Yau, 2015). Some methods assess the fraction of dropouts per gene, other methods, such as CIDR (Lin, Troup and Ho, 2017), take this process to the next step by imputing the dropout values. SAVER (Huang et al., 2017b) avoids trying to determine which observations are dropouts and aims to impute any poorly measured value using the gene-to-gene correlation pattern, and other features in the cell-type specific samples.

We apply URSM to two gene expression data sets from fetal brains, and obtain promising results on imputing single cell RNA-seq data and deconvolving bulk samples. With more upcoming single cell data on fetal brains, it would be of great scientific interest to apply URSM to specimen from different brain developing periods, which will aid our understanding on gene expression patterns during early brain development and their impact on many complex human disorders. In practice, the degrees of heterogeneity can vary for different tissues. For example, liver tissues may contain more homogeneous cell types. In all cases, URSM can be applied to obtain an accurate estimate of the cell type specific profile.

There are many existing bulk RNA-seq data sets for various human and non-human tissues that can be paired with different single cell data and jointly modeled using this unified framework. We also conduct simulation studies to demonstrate that as long as most cell types are consistent across the two data sources, URSM is robust to subtle mis-matched cell types.

As for computation, the bottleneck is the Gibbs sampling step, which scales linearly with *N, M, L* and *K*. In practice, we find that a few hundred Gibbs samples and 50-100 EM iterations are usually enough to obtain sensible results. In our experiment, for 100 single cells and 150 bulk samples, one EM iteration with 150 Gibbs samples takes about 3 minutes for 200 genes and 12 minutes for 1,000 genes using a single core on a computer equipped with an AMD Opteron(tm) Processor 6320 @ 2.8 GHz. It is straightforward to further reduce the computation time by utilizing the conditional independency to parallelize the Gibbs sampling procedure.

Many downstream analyses can be conducted with this framework. In particular, URSM provides accurate estimates of the cell type specific profile matrix, which can be used for differential expression analysis between diseased and control samples. One can also apply URSM to single cells sequenced at different developmental periods to study the developmental trajectories of the cellular profiles.

As technologies improve and costs decline, single cell analysis can move to the new level by incorporating differential expression by maternal or paternal source of the chromosome. Such information can be captured if there are genetic differences between parents in the genes. Moreover genetic variation can affect expression of genes. Already experiments are being performed to determine which genetic variants are associated with changes in single cell expression. This would allow analysis of expression based on parental origin of each copy of the gene. These sources of variation are ignored in our model. Refining and extending scRNA-seq analytical tools to accommodate these sources of variation is one of the challenges for the future.

In this paper, we present our model assuming a given number of cell types *K*. In the situation where *K* is not known a priori, one can first run the model using a larger value of *K*, examine the clustering of single cells after imputation, and then reduce to a reasonable choice of *K* by combining cells from similar clusters.

Finally, we point out that the current model is developed under the setting of supervised learning where the labels for single cells are known. One can extend this framework to conduct unsupervised cell clustering by introducing extra latent variables for cell labels in the hierarchical model. In addition, by the nature of the Multinomial distribution, the current model is fully determined by its first moment. Therefore, the imputation of single cell data may be further improved by introducing gene-gene correlations to the model. We leave the exploration in these directions to future work.

## Acknowledgements

We thank the anonymous reviewers and editor for their constructive comments and suggestions. This work was supported by SF402281, SFARI124827, R37MH057881 (Kathryn Roeder and Bernie Devlin), and R01MH109900 (Kathryn Roeder), as well as DMS-1553884 and DMS-1407771 (Jing Lei).

## APPENDIX A: DETAILS OF GIBBS SAMPLING

Recall that the latent variables for bulk data and single cell data are conditionally independent given observed data *X, Y* and parameters, so we discuss the Gibbs sampling algorithms for the two parts separately.

## A.1. Gibbs sampling for bulk data

Recall that the bulk data part of the model is equivalent to the following mixture of multinomials:

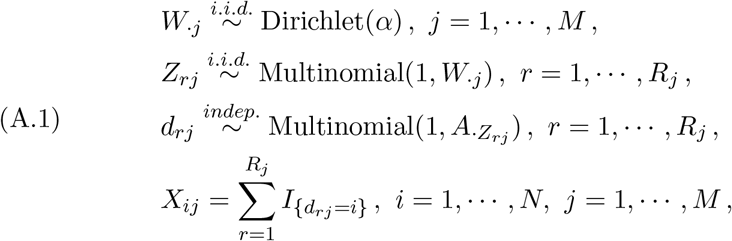

where *Z* and *d* are represented in scalars, i.e., 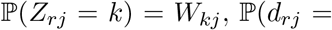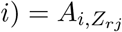. We further define

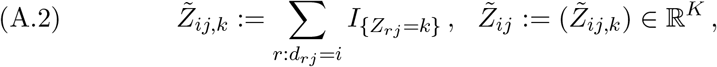

then the complete likelihood function for bulk data can be written as below:

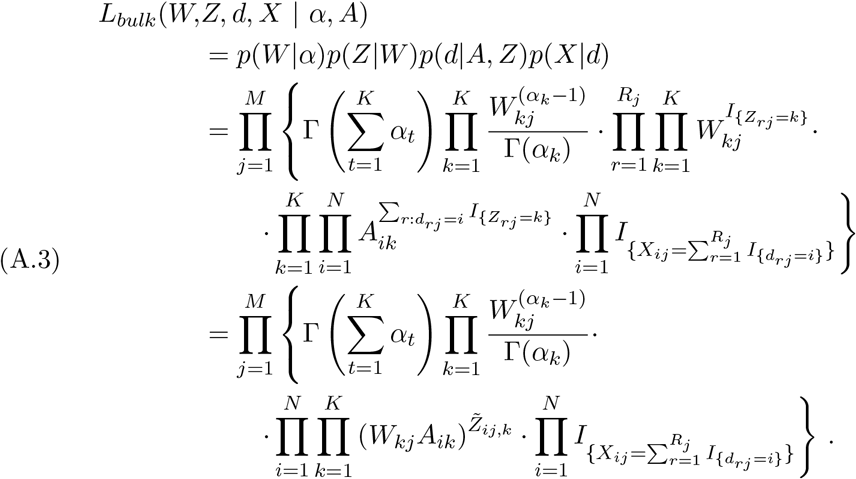

Therefore,

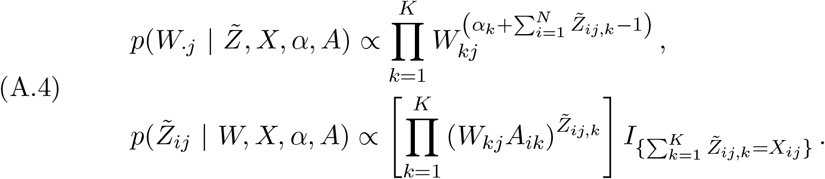

and we immediately have

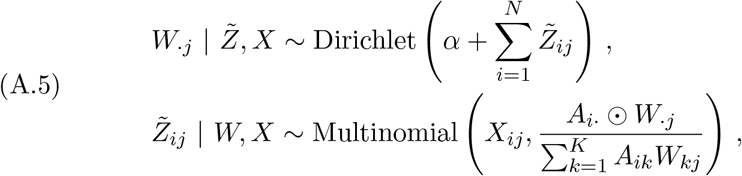

where ⨀ denotes element-wise multiplication.

## A.2. Gibbs sampling for single cell data

Recall that the first part of the single cell model can be rewritten as a Bayesian Logistic regression as follows:

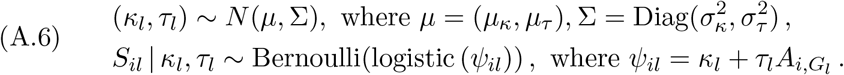

Therefore, we can utilize the data augmentation trick following Polson, Scott and Windle (2013). The key is to notice that the logistic function can be written as mixtures of Gaussians with respect to a Polya-Gamma (PG) distribution:

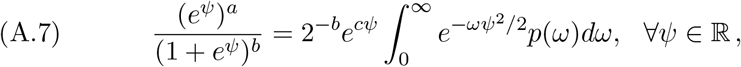

for any constants *a,b* > 0, where *c* = *a − b*/2 and *ω* ~ *PG*(0,1). Plugging in this equation, and let

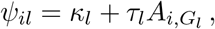

the complete likelihood for the single cell data can be written as

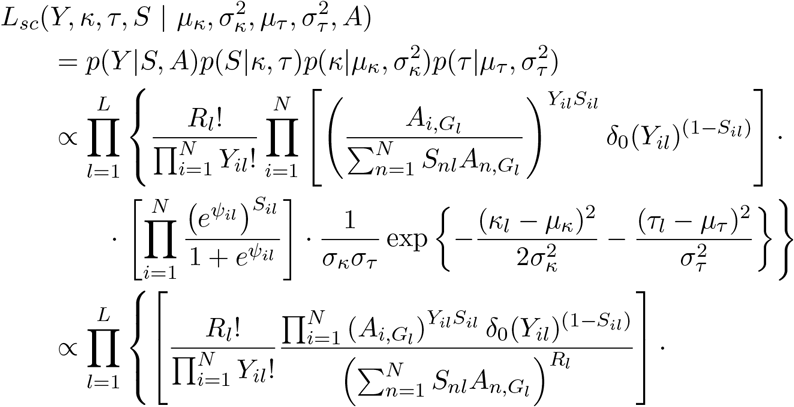

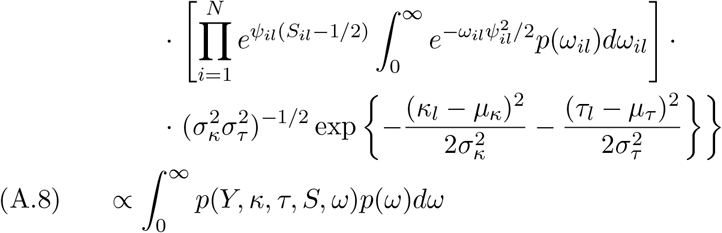

where *δ*_0_(*y*) = *I*_{*y*=0}_, and *ω_il_* ~ *PG*(0,1) independently. Then following the same arguments as in Polson, Scott and Windle (2013), we get the conditional posterior distribution for *ω_il_*, *κ_l_*, *τ_l_* as follows:

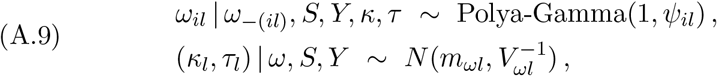

where

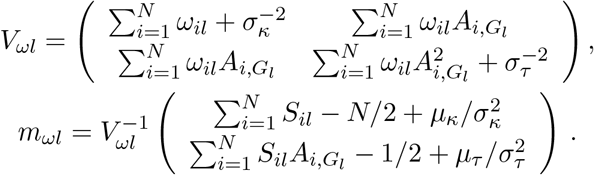

The only thing left is the conditional posterior for *S_il_*. This can be easily obtained by looking at the un-augmented version of likelihood. Note that *S_il_* is binary, and we have

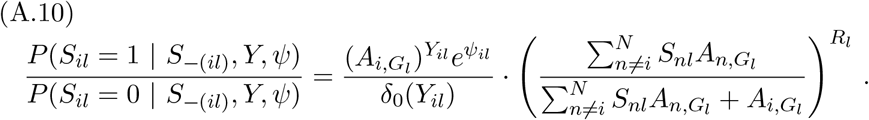

Therefore,

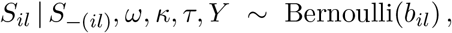

where

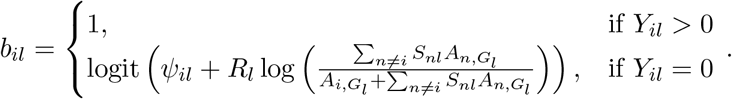

## APPENDIX B: DETAILS OF THE EM ALGORITHM

## B.1. M-step in EM algorithm

Here we give more details about the M-step in the Gibbs-EM (GEM) algorithm. By combining equations (A.3) and (A.8), the expectation of complete log likelihood function can be easily derived. However, the term of 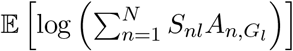 makes the optimiza tion complicated. We work around this issue by optimizing a *lower bound* of the objective function. The key step is the following lower bound using Jensen’s inequality (Paisley, 2010):

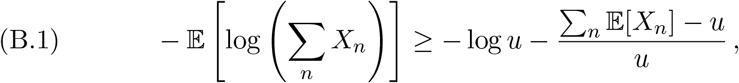

where 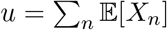, for any random variables *X_n_*’s. Using inequality (B.1), we get the following lower bound of the expected complete log likelihood, using the augmented version:

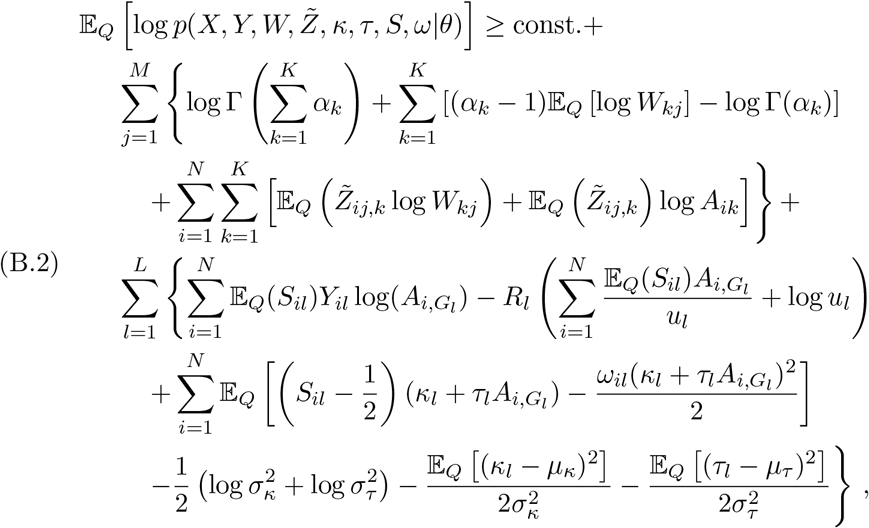

where 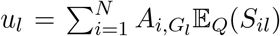, and 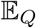 is the expectation of the posterior distribution, which can be estimated using Gibbs samples. We omit the constants that only involve the data (*X, Y*), since the goal here is to optimize over the parameters *θ* = (*A*, *α*, *μ_κ_*, 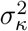, *μ_τ_*, 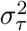). In addition, we use the fact that

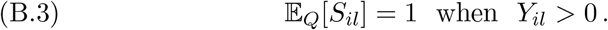

The above lower bound is also referred to as the Evidence Lower BOund (ELBO). With (B.2), it is straightforward to derive the derivatives mentioned in equation (3.9) in the original manuscript.

The final missing piece is the projection functions to the feasible set. The projection function for *α* is straightforward: for any constant *ϵ_α_* > 0,

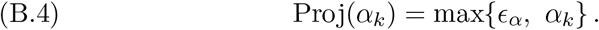

As for the profile matrix *A*, the projection function is to project onto a subset of simplex

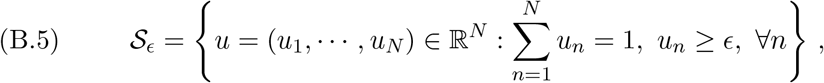

for some constant *ϵ* > 0. Wang and Carreira-Perpinan (2013) propose an efficient algorithm for the case when *ϵ* = 0. The algorithm can be easily generalized to handle a general *ϵ*, which is specified below:

*Projection algorithm for A*. For any vector *v* ∈ ℝ^*N*^, constant *ϵ* ≥ 0,

1. Sort *v* into *ṽ*, such that *ṽ*_1_ ≥ *ṽ*_2_ ≥ … ≥ *ṽ*_*N*_.
2. Find 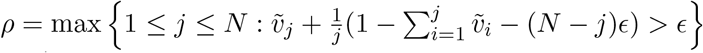.
3. Let 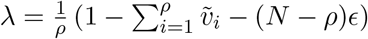.
4. Let 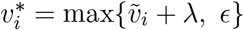, then 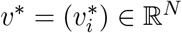 is the projection.

## B.2. Starting values

Here we present some heuristic for choosing the starting values for the EM algorithm. For the profile matrix *A*, a good candidate is the sample means *Â^naive^* in single cell data:

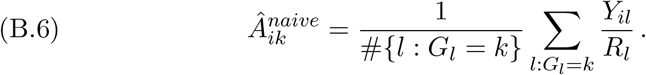

As illustrated in Figure 2b of the original manuscript, although being biased, *Â^naive^* is usually not too far away from the true profile matrix *A*.

For *α*, the starting value can be chosen using prior knowledge. Although the exact mixing proportions for each bulk sample is unknown, scientists usually have a good sense of the rough proportions of different cell types in certain tissues. In the case where prior knowledge is unavailable, α can simply be set to **1**^*K*^, which corresponds to a uniform distribution. In fact, all the simulations in the original manuscript use α_0_ = (1,1,1) as the starting value, and the performances are satisfactory.

Finally, the starting values of *μ_κ_* and *μ_τ_* can be set according to 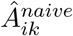, such that the distribution of 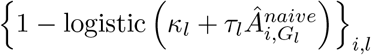 matches our prior knowledge of the dropout probabilities. In our simulation studies, *μ_κ_* is initialized at *μ*_*κ*,0_ = logit(0.4), so that the maximal dropout probability is 60% (achieved when *A_il_* = 0), and *μ_τ_* is initialized at *μ*_*τ*,0_ = (logit(0.7) − *μ*_*κ*,0_)/*Ā^naive^* where 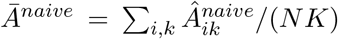. Therefore, using the initial values, the empirical average dropout probabilities is 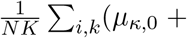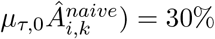.

## B.3. A MAP approximation

The bottleneck of the computation is the Gibbs sampling step in the EM algorithm. Here, we present a fast algorithm that avoids the Gibbs sampling for bulk samples by plugging in the maximum a posteriori (MAP) estimation of W. This algorithm is useful when dealing with larger data sets where computation load becomes a major concern. The simulation results in Section 4.5 of the original manuscript are obtained using this MAP approximation.

Specifically, we further assume that the sequencing depth of bulk samples is generated from *R_j_* ~ Poisson(*λ*). Then the distribution of *X* after marginalizing out *R_j_* becomes

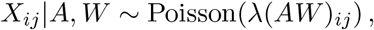

which gives

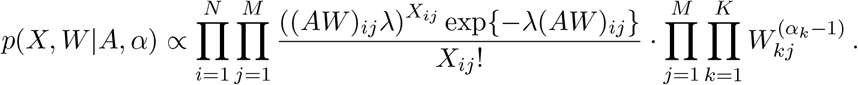

Therefore, the MAP of *W* can be obtained by

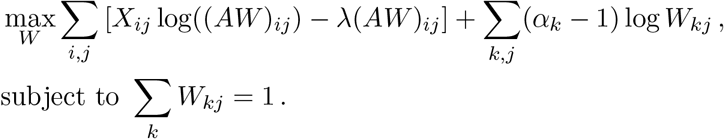

To get the optimal *W_kj_*, we write down the KKT conditions:

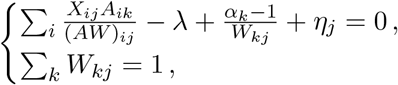

where *η_j_* is the Lagrangian multiplier for the constraint Σ*_k_ W_kj_* = 1. Finally, we obtain the following fixed-point iteration for *W*:

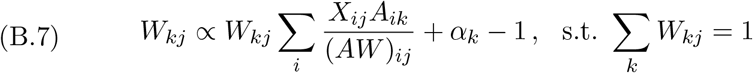

In our implementation of this fast algorithm, we perform one update of equation (B.7) in each E-step for bulk samples, starting from the *W* obtained from the previous EM iteration. Interestingly, if *α_k_* = 1, this algorithm recovers the multiplicative updates for non-negative matrix factorization when the divergence loss is used (Lee and Seung, 2001).

## REFERENCES

Abbas, A. R., Wolslegel, K., Seshasayee, D., Modrusan, Z. and Clark, H. F. (2009). Deconvolution of blood microarray data identifies cellular activation patterns in systemic lupus erythematosus. PLoS One 4 e6098.

Blei, D. M., Kucukelbir, a. and McAuliffe, J. D. (2016). Variational inference: A review for statisticians. arXiv preprint arXiv:1601.00670.

Blei, D. M., Ng, a. Y. and Jordan, M. I. (2003). Latent dirichlet allocation. Journal of machine Learning research 3 993–1022.

Brennecke, P., Anders, s., Kim, J. K., Kolodziejczyk, A. A., Zhang, X., Pros-erpio, V., Baying, B., Benes, V., Teichmann, S. A., Marioni, J. C. et al. (2013). Accounting for technical noise in single-cell RNA-seq experiments. Nature methods 10 1093–1095.

Camp, J. G., Badsha, F., Florio, M., Kanton, S., Gerber, T., Wilsch-BrÄuninger, M., Lewitus, E., Sykes, A., Hevers, W., Lancaster, M. et al. (2015). Human cerebral organoids recapitulate gene expression programs of fetal neocortex development. Proceedings of the National Academy of Sciences 112 15672–15677.

Casella, G. and George, E. I. (1992). Explaining the Gibbs Sampler. The American Statistician 46 167–174.

GTEx Consortium (2013). The Genotype-Tissue Expression (GTEx) project. Nat Genet 45 580–5.

Dempster, A. P., Laird, N. M. and Rubin, D. B. (1977). Maximum Likelihood from Incomplete Data via the EM Algorithm. JOURNAL OF THE ROYAL STATISTICAL SOCIETY, SERIES B 39 1–38.

Donoho, D. and Stodden, V. (2003). When does non-negative matrix factorization give a correct decomposition into parts? In Advances in neural information processing systems.

Dupuy, C. and Bach, F. (2016). Online but Accurate Inference for Latent Variable Models with Local Gibbs Sampling. Journal of Machine Learning Research 1.

Fan, H. C., Fu, G. K. and Fodor, S. P. (2015). Combinatorial labeling of single cells for gene expression cytometry. Science 347 1258367.

Finak, G., McDavid, A., Yajima, M., Deng, J., Gersuk, V., Shalek, A. K., Slichter, C. K., Miller, H. W., McElrath, M. J., Prlic, M., Linsley, P. S. and Gottardo, R. (2015). MAST: a flexible statistical framework for assessing transcriptional changes and characterizing heterogeneity in single-cell RNA sequencing data. Genome Biol 16 278.

Fromer, M., Roussos, P., Sieberts, S. K., Johnson, J. S., Kavanagh, D. H., Perumal, T. M., Ruderfer, D. M., Oh, E. C., Topol, A., Shah, H. R., Klei, L. L., Kramer, R., Pinto, D., Gümüs, Z. H., Cicek, A. E., Dang, K. K., Browne, A., Lu, C., Xie, L., Readhead, B., Stahl, E. A., Xiao, J., Parvizi, M., Hamamsy, T., Fullard, J. F., Wang, Y.-C., Mahajan, M. C., Derry, J. M. J., Dudley, J. T., Hemby, S. E., Logsdon, B. A., Talbot, K., Raj, T., Bennett, D. A., De Jager, P. L., Zhu, J., Zhang, B., Sullivan, P. F., Chess, A., Purcell, S. M., Shinobu, L. A., Mangrayite, L. M., Toyoshiba, H., Gur, R. E., Hahn, C.-G., Lewis, D. A., Haroutunian, V., Peters, M. A., Lipska, B. K., Buxbaum, J. D., Schadt, E. E., Hirai, K., Roeder, K., Brennand, K. J., Katsa-Nis, N., Domenici, E., Deylin, B. and Sklar, p. (2016). Gene expression elucidates functional impact of polygenic risk for schizophrenia. Nat Neurosci 19 1442–1453.

Gaujoux, R. and Seoighe, C. (2012). Semi-supervised Nonnegative Matrix Factorization for gene expression deconvolution: a case study. Infection, Genetics and Evolution 12 913–921.

Gelfand, A. and Smith, A. (1990). Sampling-Based Approaches to Calculating Marginal Densities. Journal of the American Statistical Association 85 398–409.

Geman, S. and Geman, D. (1984). Stochastic Relaxation, Gibbs Distributions, and the Bayesian Restoration of Images. IEEE Trans. Pattern Anal. Mach. Intell. 6 721–741.

Griffiths, T. L. and Steyyers, M. (2004). Finding scientific topics. Proceedings of the National academy of Sciences 101 5228–5235.

Grun, D., Lyubimoya, A., Kester, L., Wiebrands, K., Basak, O., Sasaki, N., Cleyers, H. and Van Oudenaarden, A. (2015). Single-cell messenger RNA sequencing reveals rare intestinal cell types. Nature 525 251–255.

Haque, A., Engel, J., Teichmann, S. A. and Lonnberg, T. (2017). A practical guide to single-cell RNA-sequencing for biomedical research and clinical applications. Genome Med 9 75.

Huang, W.-C., Ferris, E., Cheng, T., Horndli, C. S., Gleason, K., Tamminga, C., Wagner, J. D., Boucher, K. M., Christian, J. L. and Gregg, C. (2017a). Diverse Non-genetic, Allele-Specific Expression Effects Shape Genetic Architecture at the Cellular Level in the Mammalian Brain. Neuron 93 1094–1109.e7.

Huang, M., Wang, J., Torre, E., Dueck, H., Shaffer, S., Bonasio, R., Murray, J., Raj, A., Li, M. and Zhang, N. R. (2017b). Gene Expression Recovery For Single Cell RNA Sequencing. bioRxiv.

Jordan, M. I., Ghahramani, Z., Jaakkola, T. S. and Saul, L. K. (1999). An introduction to variational methods for graphical models. Machine learning 37 183–233.

Kang, H. J., Kawasawa, Y. I., Cheng, F., Zhu, Y., Xu, X., Li, M., Sousa, A. M., Pletikos, M., Meyer, K. A., Sedmak, G. et al. (2011). Spatio-temporal transcriptome of the human brain. Nature 478 483–489.

Kharchenko, p. V., Silberstein, L. and Scadden, D. T. (2014). Bayesian approach to single-cell differential expression analysis. Nature methods 11 740–742.

Kolodziejczyk, A. A., Kim, J. K., Svensson, V., Marioni, J. C. and Teichmann, S. A. (2015). The technology and biology of single-cell RNA sequencing. Molecular cell 58 610–620.

Lee, D. D. and Seung, H. S. (2001). Algorithms for non-negative matrix factorization. In Advances in neural information processing systems 556–562.

Lin, P., Troup, M. and Ho, J. W. K. (2017). CIDR: Ultrafast and accurate clustering through imputation for single-cell RNA-seq data. Genome Biology 18 59.

Newman, A. M., Liu, C. L., Green, M. R., Gentles, A. J., Feng, W., Xu, Y., Hoang, C. D., Diehn, M. and Alizadeh, A. A. (2015). Robust enumeration of cell subsets from tissue expression profiles. Nat Meth 12 453–457.

Padovan-Merhar, O. and Raj, A. (2013). Using variability in gene expression as a tool for studying gene regulation. Wiley Interdisciplinary Reviews: Systems Biology and Medicine 5 751–759.

Paisley, J. (2010). Two Useful Bounds for Variational Inference. Technical report.

Pierson, E. and Yau, C. (2015). ZIFA: Dimensionality reduction for zero-inflated singlecell gene expression analysis. Genome biology 16 1.

Polson, N. G., Scott, J. G. and Wlndle, J. (2013). Bayesian inference for logistic models using Pólya-Gamma latent variables. Journal of the American statistical Association 108 1339–1349.

Prabhakaran, S., Azizi, E. and Peer, D. (2016). Dirichlet Process Mixture Model for Correcting Technical Variation in Single-Cell Gene Expression Data. In Proceedings of The 33rd International Conference on Machine Learning 1070–1079.

Repsilber, D., Kern, S., Telaar, A., Walzl, G., Black, G. F., Selbig, J., Parida, S. K., Kaufmann, S. H. E. and Jacobsen, M. (2010). Biomarker discovery in heterogeneous tissue samples-taking the in-silico deconfounding approach. BMC Bioinformatics 11 27.

Satija, R., Farrell, J. A., Gennert, D., Schier, A. F. and Regev, A. (2015). Spatial reconstruction of single-cell gene expression data. Nature biotechnology 33 495–502.

Shen-Orr, S. S., Tibshirani, R., Khatri, P., Bodian, D. L., Staedtler, F., Perry, N. M., Hastie, T., Sarwal, M. M., Davis, M. M. and Butte, A. J. (2010). Cell type-specific gene expression differences in complex tissues. Nat Methods 7 287–9.

Sunkin, S. M., Ng, L., Lau, C., Dolbeare, T., Gilbert, T. L., Thompson, C. L., Hawrylycz, M. and Dang, C. (2013). Allen Brain Atlas: an integrated spatio-temporal portal for exploring the central nervous system. Nucleic Acids Res 41 D996–D1008.

Trapnell, C., Cacchiarelli, D., Grimsby, J., Pokharel, P., Li, S., Morse, M., Lennon, N. J., Livak, K. J., MiKKELSEN, T. S. and Rinn, J. L. (2014). The dynamics and regulators of cell fate decisions are revealed by pseudotemporal ordering of single cells. Nature biotechnology 32 381–386.

Vallejos, C. A., Marioni, J. C. and Richardson, S. (2015). BASiCS: Bayesian Analysis of Single-Cell Sequencing Data. PLoS Comput Biol 11 e1004333.

Vallejos, C. A., Richardson, S. and Marioni, J. C. (2016). Beyond comparisons of means: understanding changes in gene expression at the single-cell level. Genome biology 17 1.

Vallejos, C. A., Risso, D., Scialdone, A., Dudoit, S. and Marioni, J. C. (2017). Normalizing single-cell RNA sequencing data: challenges and opportunities. Nat Methods 14 565–571.

Vu, T. N., Wills, Q. F., Kalari, K. R., Niu, N., Wang, L., Rantalainen, M. and Pawitan, Y. (2016). Beta-Poisson model for single-cell RNA-seq data analyses. Bioinformatics 32 2128–35.

Wainwright, M. J. and Jordan, M. I. (2008). Graphical models, exponential families, and variational inference. Foundations and Trends@ in Machine Learning 1 1–305.

Wang, W. and Carreira-PerpinAn, M. A. (2013). Projection onto the probability simplex: An efficient algorithm with a simple proof, and an application. arXiv preprint arXiv:1309.1541.

Zhong, Y., Wan, Y.-W., Pang, K., Chow, L. M. and Liu, Z. (2013). Digital sorting of complex tissues for cell type-specific gene expression profiles. BMC bioinformatics 141.

